# Rapidly processed stool swabs approximate stool microbiota profiles

**DOI:** 10.1101/524512

**Authors:** Nicholas A. Bokulich, Juan Maldonado, Dae-Wook Kang, Rosa Krajmalnik-Brown, J. Gregory Caporaso

## Abstract

Studies of the intestinal microbiome commonly utilize stool samples to measure microbial composition in the distal gut. However, collection of stool can be difficult from some subjects under certain experimental conditions. In this study we validate the use of swabs of fecal matter to approximate measurements of microbiota in stool using 16S rRNA gene Illumina amplicon sequencing, and evaluate the effects of shipping time at ambient temperatures on accuracy. Results indicate that swab samples reliably replicate stool microbiota bacterial composition, alpha diversity, and beta diversity when swabs are processed quickly (< 2 days), but sample quality quickly degrades after this period, accompanied by increased abundances of *Enterobacteriaceae*. Fresh swabs appear to be a viable alternative to stool sampling when standard collection methods are challenging, but extended exposure to ambient temperatures prior to processing threatens sample integrity.

## Introduction

The microbial communities inhabiting the human gastrointestinal tract play important roles in digestion, immune and metabolic regulation, and disease (1). Monitoring the gut microbiota is often performed to assess the impact of disease or other disturbances (2), therapeutic interventions (3), or host development (4). Measurements of microbiota composition in the distal gut commonly utilize stool samples.

Collection and transport of stool may be difficult or impossible, however, under certain conditions, e.g., due to stool consistency or if subjects are unable or unwilling to provide stool. In a study by Sinha et al., the microbial compositions of stool swabs correlated closely with stool (5); however, this study only assessed the similarity of swab microbiota to stool at two different storage times (fresh and after 4 days at room temperature). With a similar approach, Bassis and coworkers showed that collecting and immediately processing rectal swabs also approximated stool microbiota composition (6). Rectal swabs are collected by insertion of a sterile swab into the rectum; fecal swabs are collected by applying a sterile swab to freshly passed stool or toilet paper. Collection of fecal swabs represents a simpler and less disruptive approach than either stool collection or rectal swabbing, permitting its use with sensitive patients. Swab collection also simplifies sample handling and processing during collection, archiving, and DNA extraction. This facilitates sampling under busy clinical settings or by individual subjects at home.

To validate stool swabs for measurements of intestinal microbiota, stool swabs and stool samples were collected from subjects in the autism MTT study from identical stool samples, and microbiota composition and diversity were compared between sample pairs using 16S rRNAn gene amplicon sequencing and analysis in the QIIME 2 software package (7). We show that swab and stool samples exhibit highly similar microbiota profiles, provided that the swabs were received and processed within two days of collection.

## Results

An accurate measurement of intestinal microbiota composition should demonstrate a high degree of similarity to stool composition, the current gold standard method. We measured phylogenetic similarity between samples using abundance-weighted and unweighted pairwise UniFrac distance (8). We also measured paired differences in observed richness of sequence variants, phylogenetic diversity (PD) (9), and Shannon diversity and evenness to assess alpha diversity differences between swab and stool samples.

### Fresh swab microbiota resemble stool

Freshly processed (≤ 2 days) pairs of stool and swab samples collected from the same individual at the same time (paired samples) were significantly more similar to each other than to stool or swab samples collected from the same individual but collected at different times (within-subject pairs), suggesting that stool and swab samples yield similar community structures when swabs are processed quickly (Figure 1) (weighted UniFrac Mann-Whitney U = 294.5, P = 0.007; unweighted UniFrac U = 342.5, P = 0.024). Swabs experiencing longer transport times were not significantly more similar to their stool pairs than they were to within-subject pairs (P > 0.05), suggesting that shipping times longer than 2 days do not reliably represent the microbiome of stool samples frozen at the time of collection.

**Figure 1.**
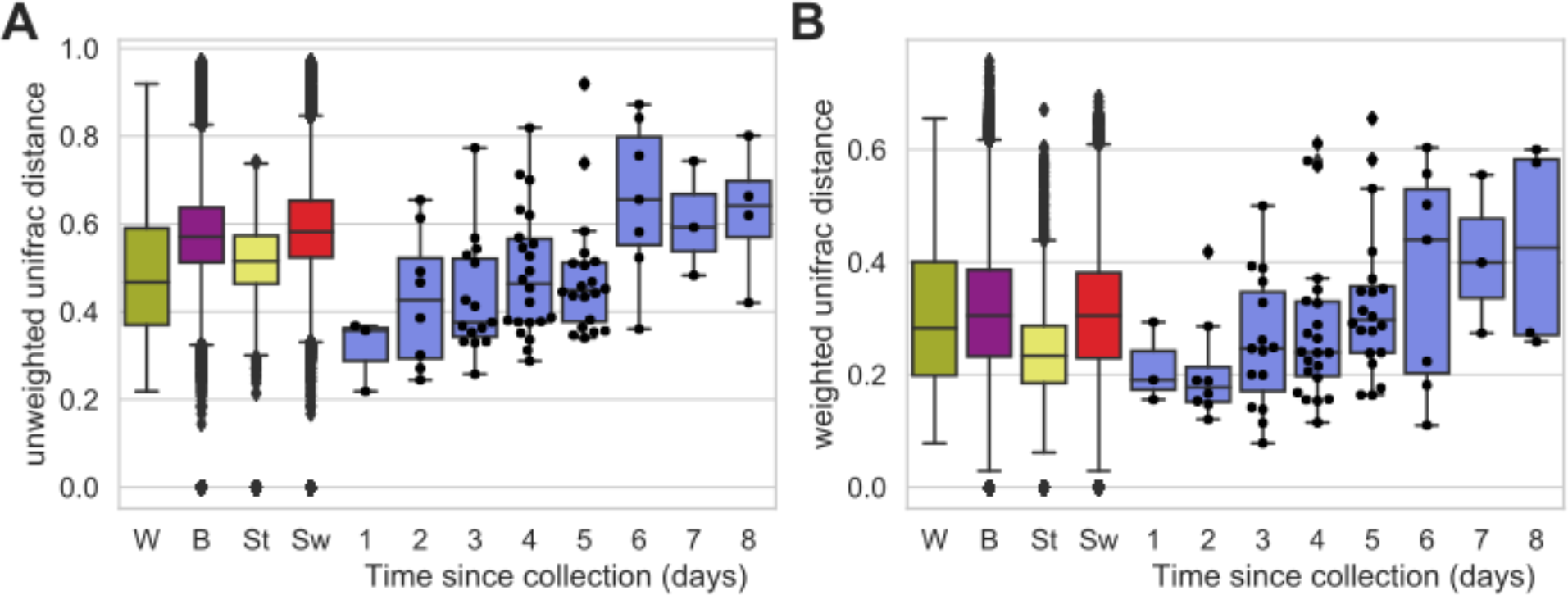
Unweighted (A) and Weighted (B) UniFrac distance distributions between sample pairs. Boxplots compare distance distributions between all samples collected from within each individual subject (“W”, green), between all subjects (“B”, purple), between all stool samples (“St”, yellow) or all swab samples (“Sw”, red) collected from the same subject at different times, and between pairs of stool and swab samples collected from the same individual at the same time (paired samples, shown in blue). Swarmplots are overlaid for paired distance measurements between swab and stool samples only, indicating the actual distribution of paired distances.

### transport time degrades swab accuracy

Both unweighted and weighted UniFrac paired sample distances increase as swab shipping time increases (Figure 1), becoming significantly more dissimilar than within-subject pairs by 6 days of shipping (Wilcoxon P < 0.05); transport time is positively correlated with paired sample dissimilarity for both weighted (Spearman R = 0.88, P = 0.004) and unweighted UniFrac (R = 0.88, P = 0.004). Thus, transport times above 1-2 days appear to have a damaging effect on swab compositional accuracy, similar to the negative effects of room-temperature storage on stool compositional accuracy (10).

Pairwise differences in alpha diversity between paired samples (swab – stool observed diversity) indicates that swab richness decreases as transport time increases (Spearman R = −0.86, P = 0.006) and PD (R = −0.88, P = 0.004). Shannon diversity (R = −0.64, P = 0.086) and evenness (R = −0.57, P = 0.139) also decrease with increasing transport time, but the correlations are not significant (Figure 2). After 4 days of transport time, swab richness, Shannon diversity, and evenness, but not PD, are significantly lower than stool (Wilcoxon P < 0.05), but transport time under 4 days does not significantly impact these alpha diversity metrics. This decrease in richness and evenness likely indicates that growth of one or more bacterial species (facultatively aerobic enterobacteria, as results below suggest) numerically overshadows the abundance of other bacteria (e.g., strict anaerobes and slower-growing organisms). The latter organisms do not disappear from this closed system, but become less likely to detect.

**Figure 2.**
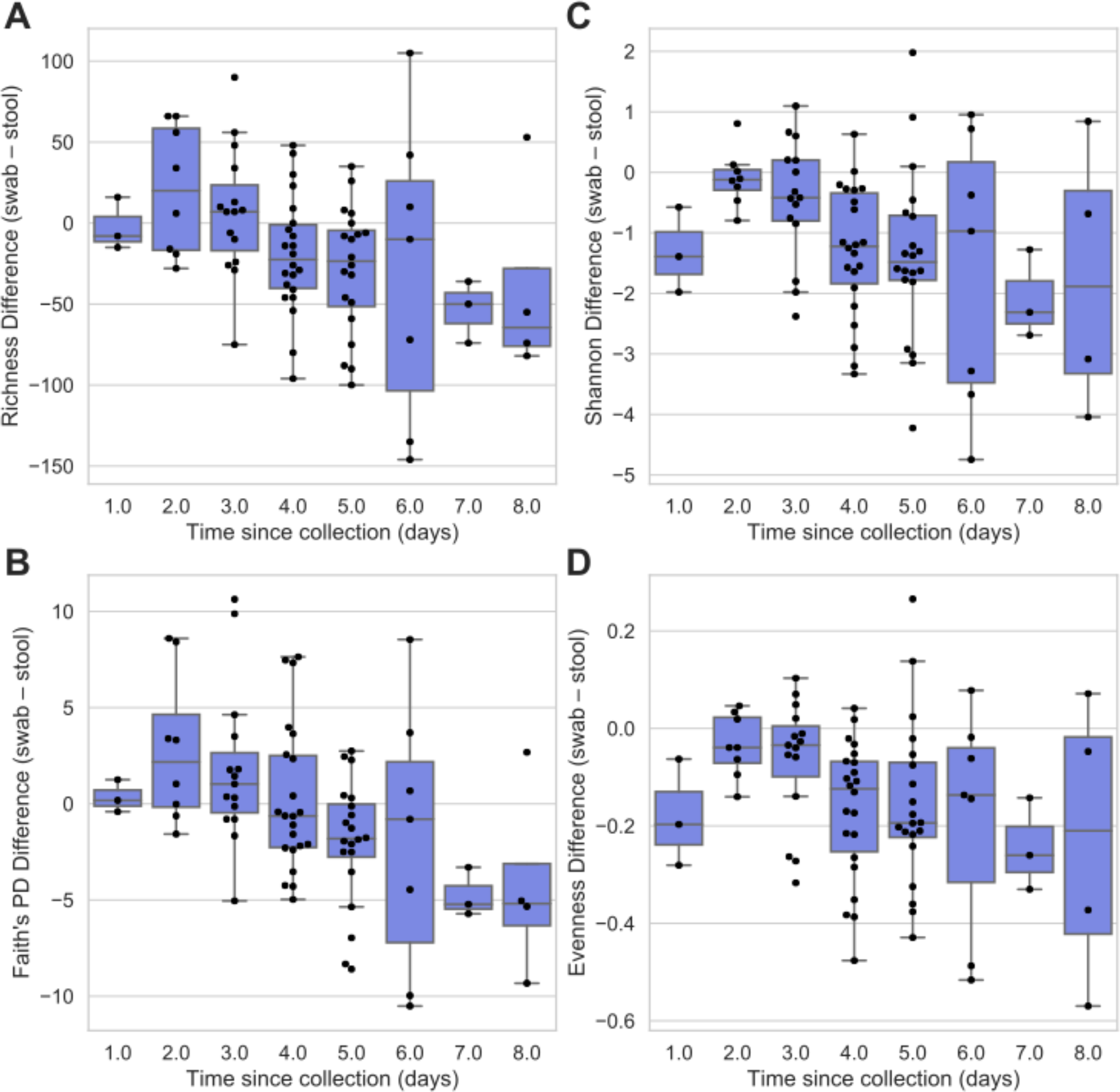
Observed differences in alpha diversity metrics between stool and swab paired samples in relation to transport time. Boxplots show quartile distributions of differences between paired samples (swab – stool observed diversity) for observed richness (A), Shannon H (B), Faith’s PD (C), and evenness (D). Swarmplots are overlaid to show actual distribution of metric differences.

### Supervised learning classification confirms accuracy of fresh swabs

To confirm the similarity of swab microbiota compared to stool microbiota, we used random forest (11) classification models to predict sample type (stool or swab) based on microbiota composition (16S rRNA gene sequence variants). Stool samples were compared to swab samples exposed to between 3-8 days of transport time (highly dissimilar from stool) or only 1-2 day of transport time (more similar to stool). Swabs exposed to 3-8 days of transport time could be accurately classified 94.6% of the time, and stool samples 90.1% of the time. However, swabs exposed to ≤ 2 days of transport time could not be reliably distinguished from stool samples: swab samples were correctly classified only 47.1% of the time (random chance is 50%). Notably, the most important features identified in each model were members of family *Enterobacteriaceae*.

### Swabs are characterized by overrepresentation of *Enterobacteriaceae* compared to stool samples

Next we determined the impact of transport time on swab bacterial taxonomic composition compared to stool to identify taxa responsible for altered diversity patterns. The taxonomic compositions of swab samples became dominated by *Enterobacteriaceae* as transport time increased, leading to a notable disparity compared to stool samples collected from the same subject at the same time (Figure 3). *Enterobacteriaceae* relative abundance was positively correlated with transport time (R = 0.88, P = 0.004) (Figure 4). Paired ANCOM tests (12) between all paired samples (regardless of transport time) indicates that bacterial species in the families *Enterobacteriaceae* and *Bacillaceae* were overrepresented in swab samples (*P* < 0.05) and a broad range of *Clostridiales* were overrepresented in stool (Table 1). While phylum Proteobacteria (represented mostly by family *Enterobacteriaceae*) was overrepresented in swab samples compared to their matching stool samples (slope > 1), most other phyla exhibited slight overrepresentation in stool (slope < 1) (Figure 5). Nevertheless, the abundances of all phyla are significantly correlated between swabs and their matching stool samples (Spearman R = 0.67, P < 0.0001) (Figure 5). This most likely indicates cellular growth of *Enterobacteriaceae* while other populations remain largely static and are supplanted at an approximately even rate. This could also indicate death and DNA degradation of these other populations, but that scenario seems much less likely given the short time frame of this experiment; however, we cannot discern changes to absolute abundance based on our compositional (relative abundance) sequence data.

**Table 1.**
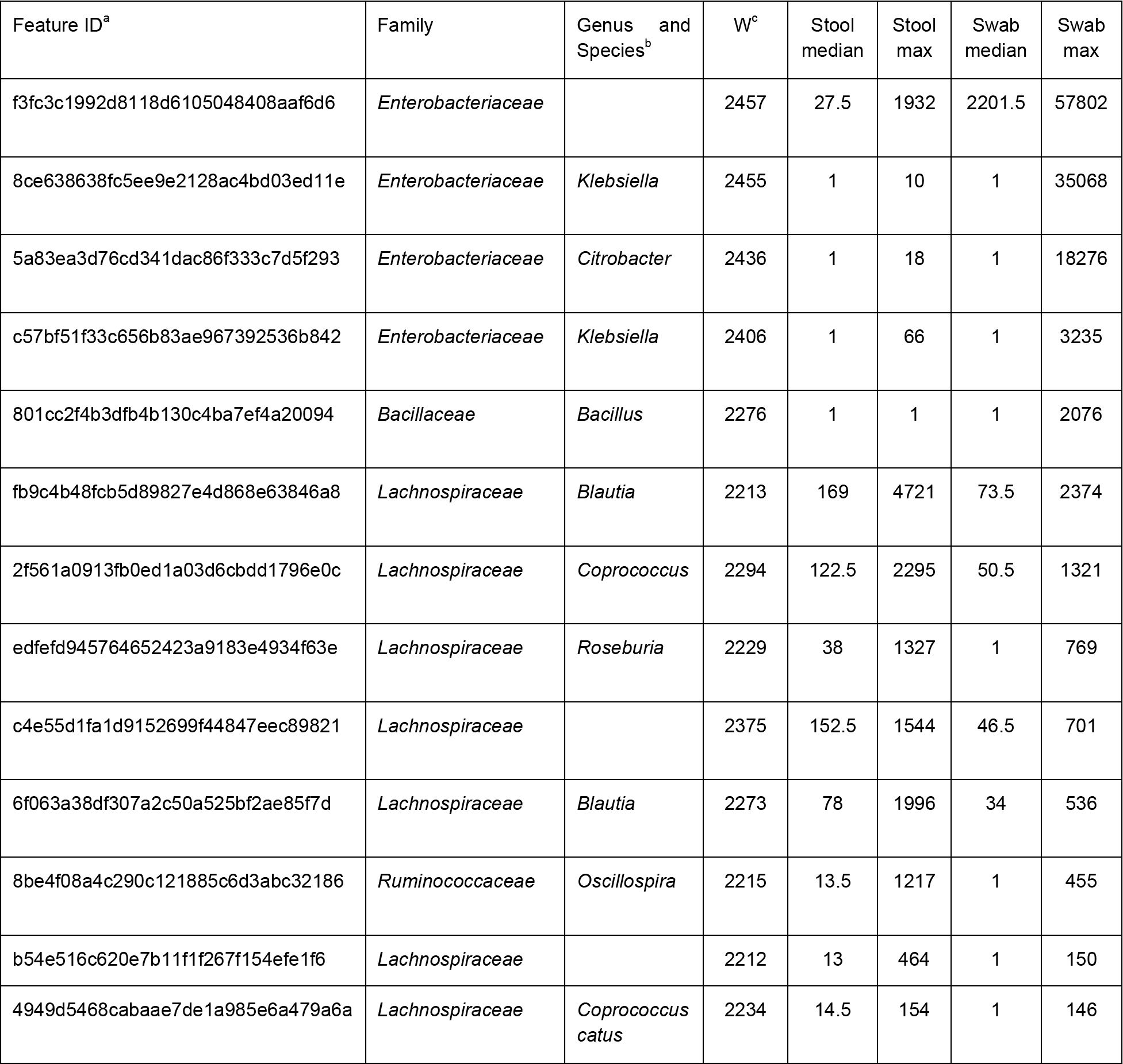
ANCOM differentially abundant sequence variants (P < 0.05) between stool and swab paired samples.

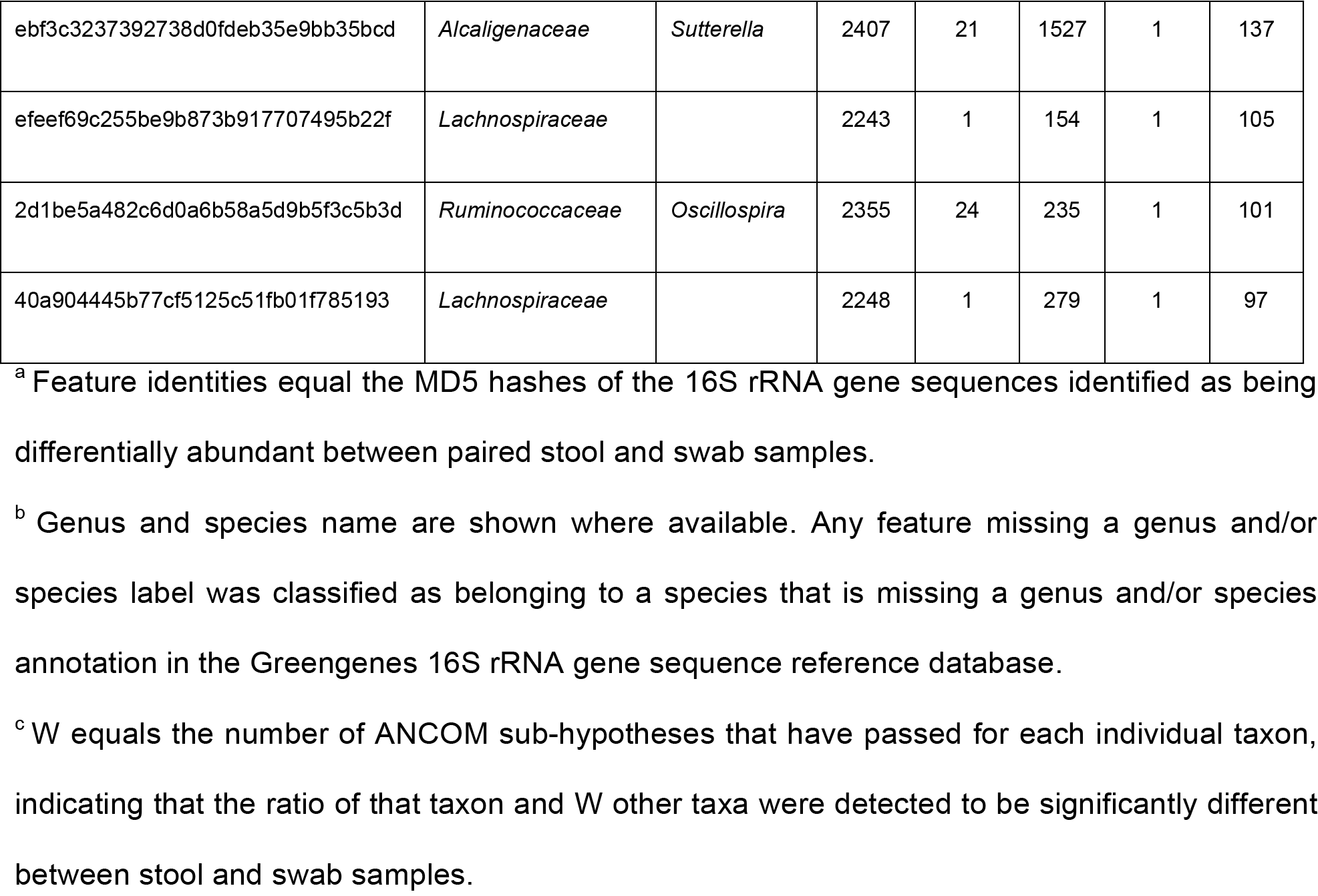

**Figure 3.**
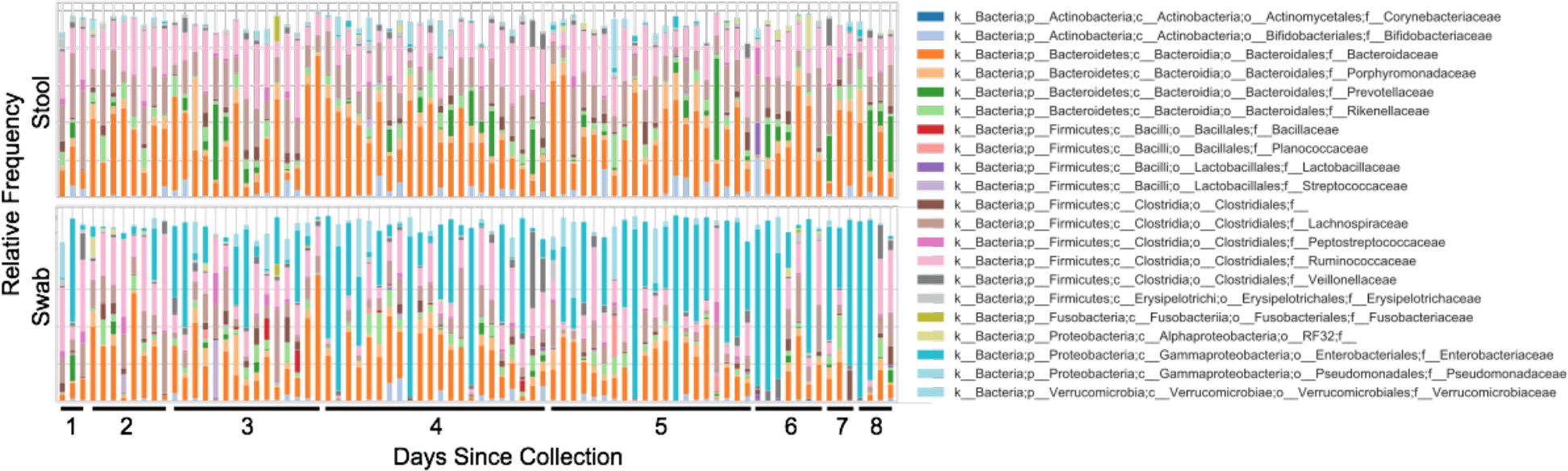
Relative abundance of bacterial families in paired stool (top) and swab samples (bottom). Paired stool and swab samples collected from the same individual at the same time point are aligned along the x axis, and sorted by swab transport time.

**Figure 4.**
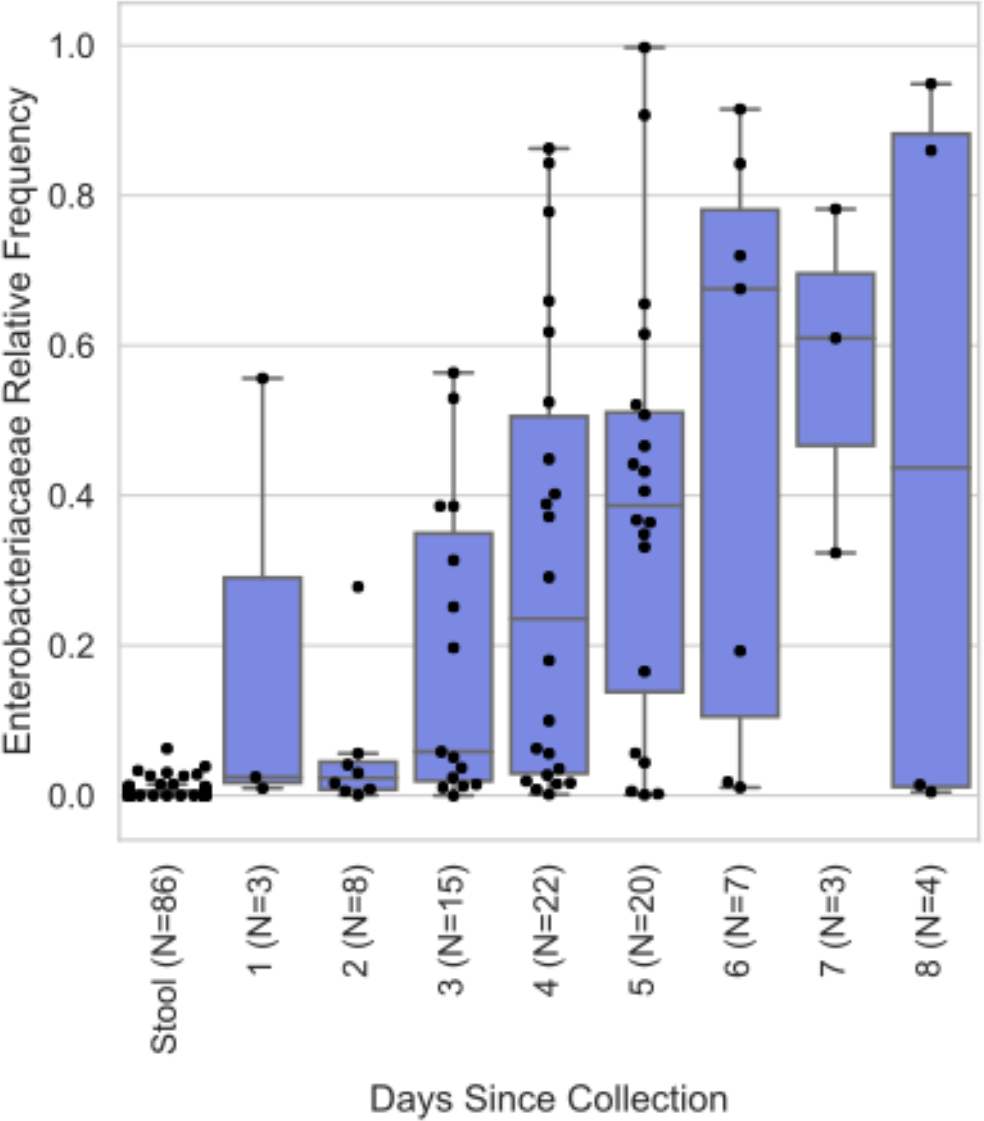
Distribution of *Enterobacteriaceae* relative frequencies in stool samples and in swab samples exposed to different transport times.

**Figure 5.**
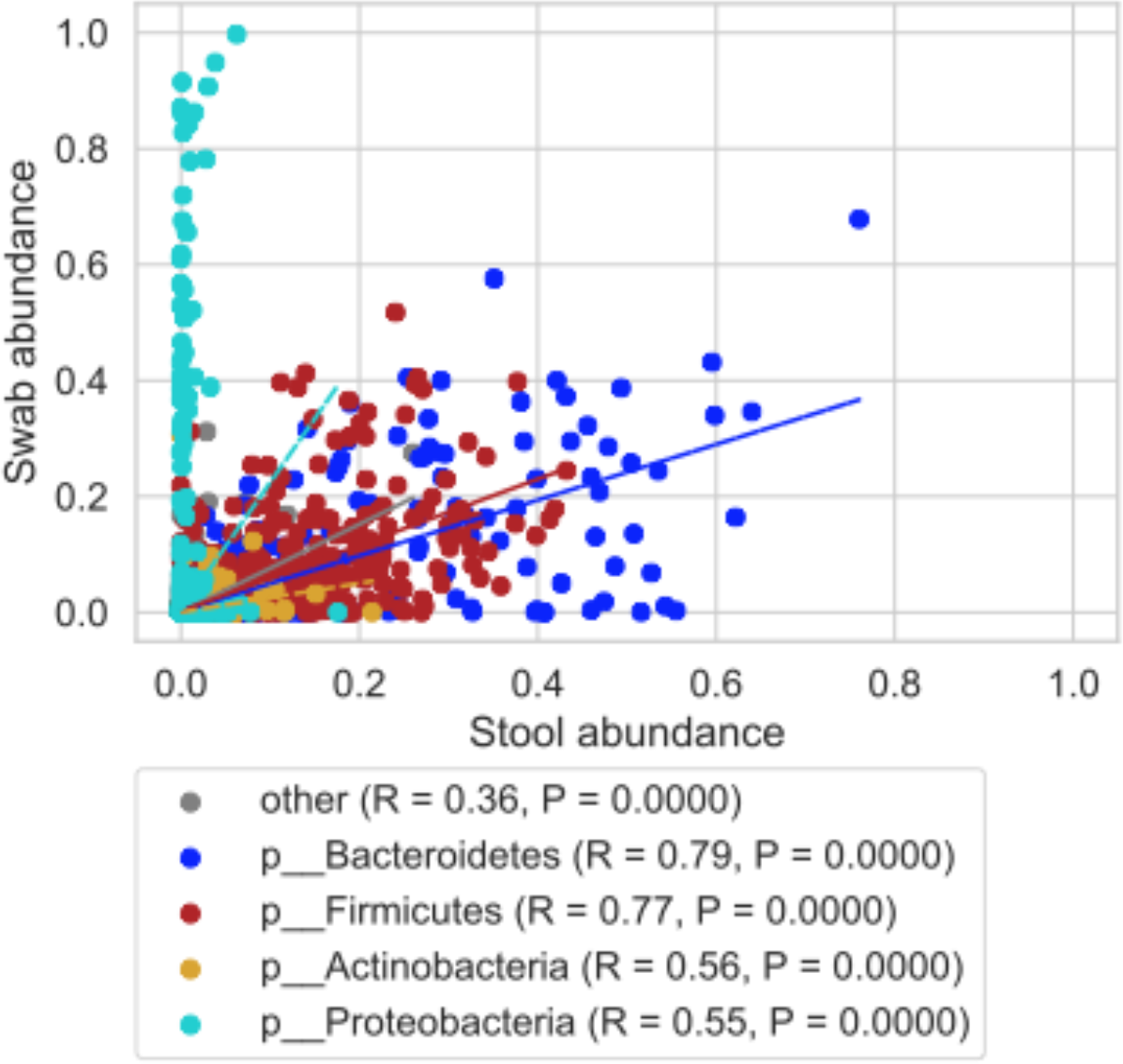
Scatterplot comparing relative abundances of all taxa observed in stool and swab samples. Taxa are colored by their phylum affiliation (all other phyla are combined into “other”), and linear regressions for each phylum are plotted. Spearman correlation coefficients (R) and P values comparing stool and swab abundances for each phylum are shown in the legend.

## Discussion

This study has demonstrated the accuracy of swabs for approximating the composition of stool samples, and evaluated the effect of transport time. Previous authors have examined the reproducibility and accuracy of fresh swabs for approximating stool microbiota measurements (5). We extend these prior studies by demonstrating the impact of storage time on swab similarity to stool. This corroborates earlier findings that swab and stool samples yield similar biological conclusions (3, 5).

We show that swabs provide an accurate approximation of stool microbiota diversity, composition, and structure, provided that the swabs are processed as freshly as possible (≤ 2 days). Stool samples and swabs could not be reliably distinguished by supervised learning classification, indicating close resemblance between these collection methods. Long transport times are associated with overrepresentation of *Enterobacteriaceae* (probably due to growth under aerobic conditions), decreasing accuracy of microbiota profiles. Prospectively, this finding could be used to further enhance the use of swabs for fecal microbiota profiling. Except in scenarios where high levels of *Enterobacteriaceae* are a normal constituent of the intestinal microbiota, such as following gastric bypass surgery (13, 14), *Enterobacteria*ceae could be used as a marker for validating swab integrity, e.g., to reject outliers that may have experienced inadequate shipping or storage; modeling compositional changes over time could also support development of algorithms to correct for biases arising from collection and storage issues.

Stool collection is not always easy or convenient. This may be due to logistical constraints (e.g., at-home collection or busy clinical settings), sample characteristics (e.g., fecal incontinence), or subject comfort. Stool swabs represent a viable alternative for measurement of distal gut microbial composition and diversity. Swabs are also considerably easier to handle and process than stool samples, streamlining collection and DNA extraction protocols. Although we find that stool and fresh swab samples could not be reliably distinguished by supervised learning classification, we do not recommend mixing stool and swab collection methods within the same study, in order to avoid introduction of experimental variation and potential sampling biases. For example, contamination and other artifactual biases could differ between collection methods and different brands of swabs, and variation should be minimized as much as possible. In studies where different collection methods become necessary, investigators should test to ensure that collection methods do not covary with other sample characteristics or metadata.

## Materials and methods

### Data availability

This study re-analyzed a previously published 16S rRNA gene sequence dataset (3), which is available in the open-source microbiome database Qiita (qiita.microbio.me) under the study ID number 10532.

### Sample collection and processing

Stool samples and swabs were collected and processed as previously described in a study of autistic children receiving microbiota transfer therapy (3). Stool samples and fecal swabs were collected by subjects’ parents. Fecal samples were stored in dry ice and collected by a driver, and frozen at −80°C immediately upon arrival at the laboratory. Swabs were shipped to the lab by standard postal mail. After defecation, fecal matter was collected from toilet paper using a sterile swab (Fisher Scientific BD Culture Swab item number B4320135), taking care not to contact the paper or overload the swab. Samples were shipped at room temperature and frozen at −80°C immediately upon arrival at the laboratory. Swab samples were primarily shipped within Arizona at different times of year, so temperatures (and hence shipping effects) may be slightly greater than other regions. The time between shipping and receipt was logged as “days in transit”, as used to perform statistical analyses described below. DNA extraction and sequencing were performed as previously described, following the earth microbiome project standard protocol for 16S V4 rRNA gene sequencing with 515f-806r primers (15). A total of 123 stools and 355 swabs were collected and analyzed in the current study, including 98 pairs of stool and swab samples that were collected from the same source stool. Swab transport times varied from 0 to 68 days; however, only days 1-8 contained sufficient sample size (minimum N = 3 stool-swab pairs) and were used for assessing the impact of transport time on swab composition accuracy compared to paired stools.

### Microbiome analysis

Sequence data were processed and analyzed using QIIME 2 (7). Raw sequences were quality-filtered using DADA2 (16) to remove PhiX, chimeric, and erroneous reads. Sequence variants were aligned using mafft (17) and used to construct a phylogenetic tree using fasttree2 (18). Taxonomy was assigned to sequence variants using q2-feature-classifier (19) against the GreenGenes 16S rRNA reference database 13_8 release (20).

### Statistical analysis

QIIME 2 was used to measure the following microbiota alpha diversity metrics: richness (as observed sequence variants), Shannon diversity and evenness, and Phylogenetic Diversity (9). Microbiome beta diversity was estimated in QIIME 2 using weighted and unweighted UniFrac distance (8). Feature tables were evenly subsampled at 5,000 sequences per sample prior to alpha or beta diversity analyses.

Alpha diversity differences and UniFrac distances between paired stool and swab samples from identical source samples (paired samples) were calculated using q2-longitudinal (21). ANCOM (12) was used to test whether the abundances of individual taxa differed between paired samples. Balance trees analysis and ordinary least squares regression on balances was performed using the q2-gneiss plugin (22). Spearman correlation coefficients were computed between transport time and median alpha diversity metrics, UniFrac distance, and *Enterobacteriaceae* relative abundance. Mann-Whitney U tests were used to test whether relative abundances of family *Enterobacteriaceae* were significantly different between stool samples and swab samples exposed to different transport times. Supervised learning classification was performed in q2-sample-classifier (23), using random forests classifiers (11) grown with 500 trees, trained on a random subset of the data (80%) and validated on the remaining samples.

## Acknowledgments

The authors thank James B. Adams (Arizona State University) for useful discussions. This work was supported in part by the Arizona Board of Regents, The Autism Research Institute, and the National Science Foundation (award 1565100 to JGC).

## References

1. Gilbert JA, Blaser MJ, Caporaso JG, Jansson JK, Lynch SV, Knight R. 2018. Current understanding of the human microbiome. Nat Med 24:392–400.

2. Gevers D, Kugathasan S, Denson LA, Vázquez-Baeza Y, Van Treuren W, Ren B, Schwager E, Knights D, Song SJ, Yassour M, Morgan XC, Kostic AD, Luo C, González A, McDonald D, Haberman Y, Walters T, Baker S, Rosh J, Stephens M, Heyman M, Markowitz J, Baldassano R, Griffiths A, Sylvester F, Mack D, Kim S, Crandall W, Hyams J, Huttenhower C, Knight R, Xavier RJ. 2014. The treatment-naive microbiome in new-onset Crohn’s disease. Cell Host Microbe 15:382–392.

3. Kang D-W, Adams JB, Gregory AC, Borody T, Chittick L, Fasano A, Khoruts A, Geis E, Maldonado J, McDonough-Means S, Pollard EL, Roux S, Sadowsky MJ, Lipson KS, Sullivan MB, Caporaso JG, Krajmalnik-Brown R. 2017. Microbiota Transfer Therapy alters gut ecosystem and improves gastrointestinal and autism symptoms: an open-label study. Microbiome 5:10.

4. Bokulich NA, Chung J, Battaglia T, Henderson N, Jay M, Li H D, Lieber A, Wu F, Perez-Perez GI, Chen Y, Schweizer W, Zheng X, Contreras M, Dominguez-Bello MG, Blaser MJ. 2016. Antibiotics, birth mode, and diet shape microbiome maturation during early life. Sci Transl Med 8:343ra82.

5. Sinha R, Chen J, Amir A, Vogtmann E, Shi J, Inman KS, Flores R, Sampson J, Knight R, Chia N. 2016. Collecting Fecal Samples for Microbiome Analyses in Epidemiology Studies. Cancer Epidemiol Biomarkers Prev 25:407–416.

6. Bassis CM, Moore NM, Lolans K, Seekatz AM, Weinstein RA, Young VB, Hayden MK, CDC Prevention Epicenters Program. 2017. Comparison of stool versus rectal swab samples and storage conditions on bacterial community profiles. BMC Microbiol 17:78.

7. Bolyen E, Rideout JR, Dillon MR, Bokulich NA, Abnet C, Al-Ghalith GA, Alexander H, Alm EJ, Arumugam M, Asnicar F, Bai Y, Bisanz JE, Bittinger K, Brejnrod A, Brislawn CJ, Titus Brown C, Callahan BJ, Caraballo-Rodríguez AM, Chase J, Cope E, Da Silva R, Dorrestein PC, Douglas GM, Durall DM, Duvallet C, Edwardson CF, Ernst M, Estaki M, Fouquier J, Gauglitz JM, Gibson DL, Gonzalez A, Gorlick K, Guo J, Hillmann B, Holmes S, Holste H, Huttenhower C, Huttley G, Janssen S, Jarmusch AK, Jiang L, Kaehler B, Kang KB, Keefe CR, Keim P, Kelley ST, Knights D, Koester I, Kosciolek T, Kreps J, Langille MGI, Lee J, Ley R, Liu Y-X, Loftfield E, Lozupone C, Maher M, Marotz C, Martin BD, McDonald D, McIver LJ, Melnik AV, Metcalf JL, Morgan SC, Morton J, Naimey AT, Navas-Molina JA, Nothias LF, Orchanian SB, Pearson T, Peoples SL, Petras D, Preuss ML, Pruesse E, Rasmussen LB, Rivers A, Michael S Robeson II, Rosenthal P, Segata N, Shaffer M, Shiffer A, Sinha R, Song SJ, Spear JR, Swafford AD, Thompson LR, Torres PJ, Trinh P, Tripathi A, Turnbaugh PJ, Ul-Hasan S, van der Hooft JJJ, Vargas F, Vázquez-Baeza Y, Vogtmann E, von Hippel M, Walters W, Wan Y, Wang M, Warren J, Weber KC, Williamson CHD, Willis AD, Xu ZZ, Zaneveld JR, Zhang Y, Knight R, Gregory Caporaso J. 2018. QIIME 2: Reproducible, interactive, scalable, and extensible microbiome data science e27295v1. PeerJ Preprints.

8. Lozupone C, Knight R. 2005. UniFrac: a new phylogenetic method for comparing microbial communities. Appl Environ Microbiol 71:8228–8235.

9. Faith DP. 1992. Conservation evaluation and phylogenetic diversity. Biol Conserv 61:1–10.

10. Song SJ, Amir A, Metcalf JL, Amato KR, Xu ZZ, Humphrey G, Knight R. 2016. Preservation Methods Differ in Fecal Microbiome Stability, Affecting Suitability for Field Studies. mSystems 1.

11. Breiman L. 2001. Random forests. Mach Learn 45:5–32.

12. Mandal S, Van Treuren W, White RA, Eggesbø M, Knight R, Peddada SD. 2015. Analysis of composition of microbiomes: a novel method for studying microbial composition. Microb Ecol Health Dis 26:27663.

13. Zhang H, DiBaise JK, Zuccolo A, Kudrna D, Braidotti M, Yu Y, Parameswaran P, Crowell MD, Wing R, Rittmann BE, Krajmalnik-Brown R. 2009. Human gut microbiota in obesity and after gastric bypass. Proceedings of the National Academy of Sciences 106:2365–2370.

14. Ilhan ZE, DiBaise JK, Isern NG, Hoyt DW, Marcus AK, Kang D-W, Crowell MD, Rittmann BE, Krajmalnik-Brown R. 2017. Distinctive microbiomes and metabolites linked with weight loss after gastric bypass, but not gastric banding. ISME J 11:2047–2058.

15. Caporaso JG, Lauber CL, Walters WA, Berg-Lyons D, Huntley J, Fierer N, Owens SM, Betley J, Fraser L, Bauer M, Gormley N, Gilbert JA, Smith G, Knight R. 2012. Ultra-high-throughput microbial community analysis on the Illumina HiSeq and MiSeq platforms. ISME J 6:1621–1624.

16. Callahan BJ, McMurdie PJ, Rosen MJ, Han AW, Johnson AJA, Holmes SP. 2016. DADA 2: High-resolution sample inference from Illumina amplicon data. Nat Methods 13:581–583.

17. Katoh K, Misawa K, Kuma K-I, Miyata T. 2002. MAFFT: a novel method for rapid multiple sequence alignment based on fast Fourier transform. Nucleic Acids Res 30:3059–3066.

18. Price MN, Dehal PS, Arkin AP. 2010. FastTree 2--approximately maximum-likelihood trees for large alignments. PLoS One 5:e9490.

19. Bokulich NA, Kaehler BD, Rideout JR, Dillon M, Bolyen E, Knight R, Huttley GA, Gregory Caporaso J. 2018. Optimizing taxonomic classification of marker-gene amplicon sequences with QIIME 2’s q2-feature-classifier plugin. Microbiome 6:90.

20. McDonald D, Price MN, Goodrich J, Nawrocki EP, DeSantis TZ, Probst A, Andersen GL, Knight R, Hugenholtz P. 2012. An improved Greengenes taxonomy with explicit ranks for ecological and evolutionary analyses of bacteria and archaea. ISME J 6:610–618.

21. Bokulich NA, Dillon MR, Zhang Y, Rideout JR, Bolyen E, Li H, Albert PS, Caporaso JG. 2018. q2-longitudinal: Longitudinal and Paired-Sample Analyses of Microbiome Data. mSystems 3:e00219–18.

22. Morton JT, Sanders J, Quinn RA, McDonald D, Gonzalez A, Vázquez-Baeza Y, Navas-Molina JA, Song SJ, Metcalf JL, Hyde ER, Lladser M, Dorrestein PC, Knight R. 2017. Balance Trees Reveal Microbial Niche Differentiation. mSystems 2.

23. Bokulich N, Dillon M, Bolyen E, Kaehler BD, Huttley GA, Gregory Caporaso J. 2018. q2-sample-classifier: machine-learning tools for microbiome classification and regression. Journal of Open Source Software 3:934.

